# In situ detection of protein interactions for recombinant therapeutic enzymes

**DOI:** 10.1101/2020.05.06.081885

**Authors:** Mojtaba Samoudi, Chih-Chung Kuo, Caressa M. Robinson, Km Shams-Ud-Doha, Song-Min Schinn, Stefan Kol, Linus Weiss, Sara Petersen Bjorn, Bjorn G. Voldborg, Alexandre Rosa Campos, Nathan E. Lewis

**Affiliations:** Dept of Pediatrics, University of California, San Diego; Novo Nordisk Foundation Center for Biosustainability at UC San Diego; Dept of Bioengineering, University of California, San Diego; Sanford Burnham Prebys Medical Discovery Institute; Novo Nordisk Foundation Center for Biosustainability, Technical University of Denmark; Dept of Biochemistry, Eberhard Karls University of Tübingen, Germany

**Keywords:** Therapeutic proteins, Secretory pathway, BioID, Cell engineering, Disulfide bond

## Abstract

Despite their therapeutic potential, many protein drugs remain inaccessible to patients since they are difficult to secrete. Each recombinant protein has unique physicochemical properties and requires different machinery for proper folding, assembly, and post-translational modifications (PTMs). Here we aimed to identify the machinery supporting recombinant protein secretion by measuring the protein-protein interaction (PPI) networks of four different recombinant proteins (SERPINA1, SERPINC1, SERPING1 and SeAP) with various PTMs and structural motifs using the proximity-dependent biotin identification (BioID) method. We identified PPIs associated with specific features of the secreted proteins using a Bayesian statistical model, and found proteins involved in protein folding, disulfide bond formation and N-glycosylation were positively correlated with the corresponding features of the four model proteins. Among others, oxidative folding enzymes showed the strongest association with disulfide bond formation, supporting their critical roles in proper folding and maintaining the ER stability. Knockdown of disulfide-isomerase PDIA4, a measured interactor with significance for SERPINC1 but not SERPINA1, led to the decreased secretion of SERPINC1, which relies on its extensive disulfide bonds, compared to SERPINA1, which has no disulfide bonds. Proximity-dependent labeling successfully identified the transient interactions supporting synthesis of secreted recombinant proteins and refined our understanding of key molecular mechanisms of the secretory pathway during recombinant protein production.

## Introduction

Therapeutic proteins are increasingly important for treating diverse diseases, including cancers, autoimmunity/inflammation, infectious diseases, and genetic disorders. For example, the plasma protein therapeutics market is expected to grow by $36 billion (USD) by 2024. Mammalian cells are the dominant production system due to their ability to perform PTMs that are required for drug safety and function (Jenkins, Murphy, & Tyther, 2008; Matasci, Hacker, Baldi, & Wurm, 2008). However, the complexities associated with the mammalian secretory machinery remains a bottleneck in recombinant protein production (Gutierrez et al., 2020). The secretory pathway machinery includes >575 gene products tasked with the synthesis, folding, PTMs, quality control, and trafficking of secreted proteins (SecPs) (Feizi, Gatto, Uhlen, & Nielsen, 2017; Lund et al., 2017; Novick, Ferro, & Schekman, 1981; Reynaud & Simpson, 2002). Numerous components of the secretory pathway (SecMs) have been engineered to increase the capacity of the secretion (Xiao, Shiloach, & Betenbaugh, 2014). However, the precision and efficiency of the mammalian secretory pathway results from the coordinated effort of these secretory machinery components including chaperones, modifying enzymes (e.g., protein disulfide isomerases and glycosyltransferases), and transporters within the secretory pathway. Overexpression of heterologous proteins in this tightly regulated and complex system could impact its functionality and homeostasis, resulting in adaptive responses that can impair both protein quantity and quality (Hussain, Maldonado-Agurto, & Dickson, 2014; Young, Yuraszeck, & Robinson, 2011). More importantly, variability in the structures and modifications of recombinant proteins could necessitate a customized secretion machinery to handle this diversity, but the secretory machinery of recombinant protein producing cells has not been adapted to facilitate the high titer secretion desired for most recombinant proteins. A previous study also showed human protein secretory pathway genes are expressed in a tissue-specific pattern to support the diversity of secreted proteins and their modifications (Feizi et al., 2017), suggesting that expression of several SecMs is regulated to support client SecPs in the secretory pathway. Unfortunately, the SecMs needed to support any given secreted protein remain unknown. Thus, there is a need to elucidate the SecMs that support the expression of different recombinant proteins with specific features. This can guide mammalian synthetic biology efforts to engineer enhanced cells capable of expressing proteins of different kinds in a client-specific manner.

PPI networks are invaluable tools for deciphering the molecular basis of biological processes. New proximity dependent labeling methods such as BioID (Kim et al., 2014; Roux, Kim, Raida, & Burke, 2012) and APEX (Rhee et al., 2013) can identify weak and transient interactions in living cells, along with stable interactions. Furthermore, BioID offers a high-throughput approach for systematic detection of intracellular PPIs occurring in various cellular compartments and has been used to characterize PPI networks and subcellular organization (Varnaitė & MacNeill, 2016). BioID relies on expressing a protein of interest fused to a promiscuous biotin ligase (BirA) that can biotinylate the proximal interactors in nanometer-scale labeling radius (Kim et al., 2014). For example, this approach has mapped protein interactions at human centrosomes and cilia (Firat-Karalar & Stearns, 2015; Gupta et al., 2015), focal adhesions (Dong et al., 2016), nuclear pore (Kim et al., 2014) and ER membrane-bound ribosomes (Hoffman, Chen, Zheng, & Nicchitta, 2019). BirA has also been used in proximity-specific ribosome profiling to reveal principles of ER co-translational translocation (Jan, Williams, & Weissman, 2014). Here we used BioID2, an improved smaller biotin ligase for BioID (Kim et al., 2016; Varnaitė & MacNeill, 2016), to explore how the SecMs involved vary for different secreted therapeutic proteins (Fig. 1). Specifically, BioID2 was employed to identify SecMs that interact with three SERPIN-family proteins (SERPINA1: treatment for Alpha-1-antitrypsin deficiency, SERPINC1: treatment for Hereditary antithrombin deficiency, and SERPING1: treatment for acute attacks of hereditary angioedema) and secreted embryonic alkaline phosphatase (SeAP), which is a truncated form of Alkaline Phosphatase, Placental Type (ALPP). These proteins vary in their PTMs (e.g., glycosylation, disulfide bond and residue modifications) and have different amino acid sequences that consequently form different local motifs. Using a Bayesian statistical model, we identified the critical PPIs that are positively correlated with each protein feature. Identification of these PPIs will refine our understanding of how the secretory pathway functions during the expression of the recombinant proteins and introduce novel targets for secretory pathway engineering in a client specific manner.

**Figure 1.**
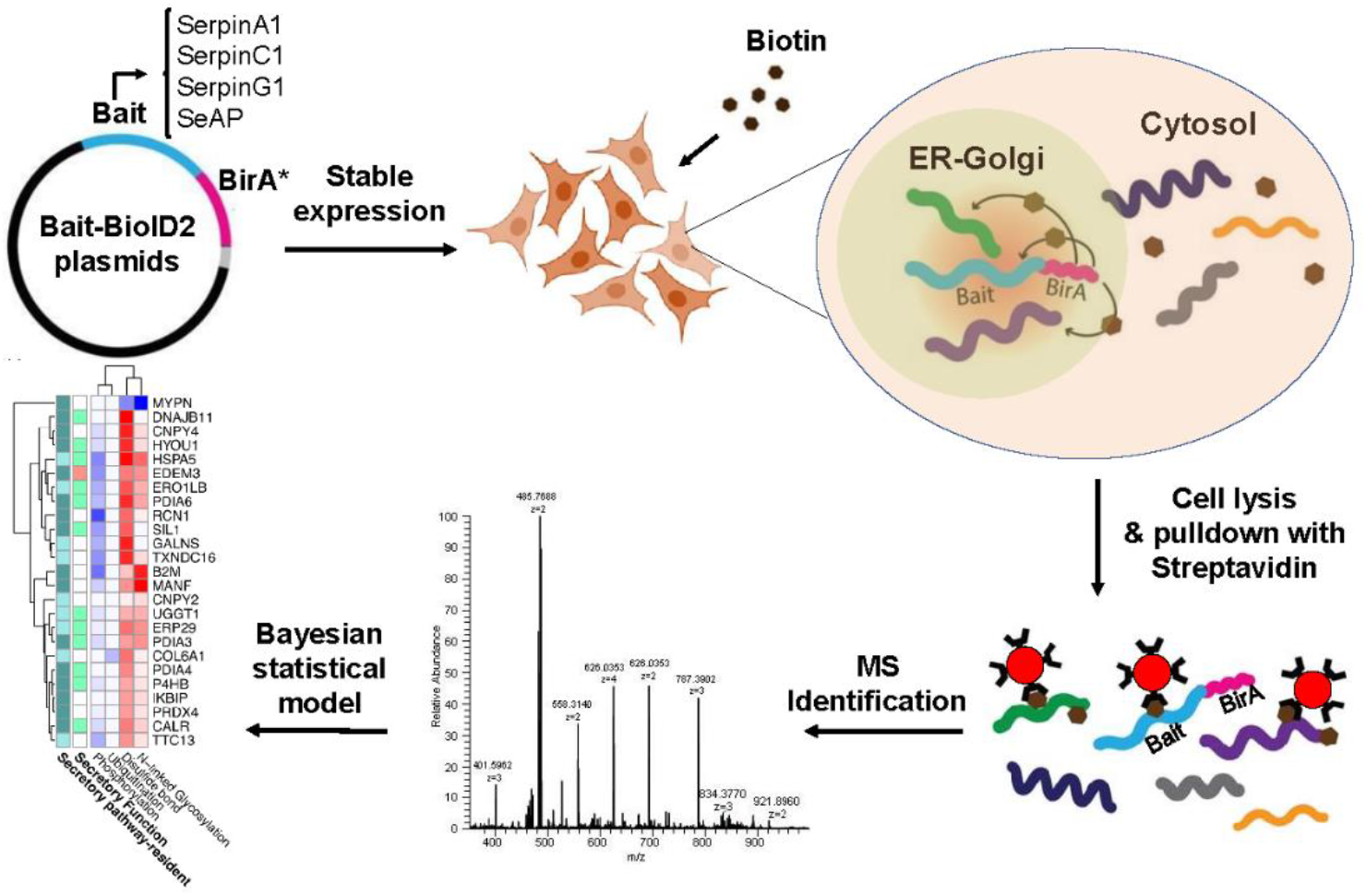
Flowchart of the BioID2 application to detect in situ interactions supporting therapeutic proteins secretion.

## Materials and Methods

### Molecular cloning and generation of stable cell lines

All plasmids used in this study were constructed by PCR and Gibson isothermal assembly. The expression ORFs, hereafter named bait-BirA, were constructed by fusing BioID2 to the C-terminal of each model protein (with a glycine-serine linker added between) and a 3XFLAG tag at C-terminal to simplify the immuno detection. ORFs were inserted into pcDNA5/FRT (Invitrogen), which allows targeted integration of the transgenes into the host genome. Gibson assembly primers were designed by SnapGene software and used to amplify the corresponding fragments and vectors with long overlapping overhangs, which were then assembled using Gibson Assembly Master Mix (NEB). To obtain secretable BioID2 (without any bait protein), Gibson assembly was employed to fuse the signal peptide of SERPINC1 gene to the N-terminal of BirA (hereafter referred to as Signal-BirA). Assembly products were transformed to the chemically competent *E. coli*, and recombinant plasmids were verified by restriction digestion and sequencing. For all experiments, Flp-In 293 cells (Invitrogen) were cultured in DMEM media supplemented with fetal bovine serum (10%) and antibiotics (penicillin, 100 U mL-1 and streptomycin, 100 μg mL-1) and maintained at 37 °C under 5 % CO2. For generating stable cell lines, Flp-In 293 cells were seeded in 6 well plates at a density of 0.5×10^6^ cells per well the day before transfection. Cells were then co-transfected with each pcDNA5/FRT vector containing expression cassette and pOG44 plasmid using Lipofectamine^®^ 2000 according to the manufacturer’s directions. After recovery from transfection, cells were grown in DMEM containing 10% FBS, 1% PenStrep, and 150 μg/mL Hygromycin B to select hygromycin-resistant cells. Individual resistant colonies were isolated, pooled, and seeded in 24-well plates for further scaling up and screened for expression of the fusion proteins by Western Blotting.

### Immunofluorescence

Recombinant HEK293 cells expressing BioID2 fusions were grown in complete medium supplemented with 50 μM biotin on coverslips until 70% confluent. Cells were then fixed in PBS containing 4% PFA for 10 min at room temperature. Blocking was performed by incubating fixed cells with 1% BSA and 5% normal goat serum in PBST. Anti-flag mouse monoclonal antibody-Dylight 650 conjugate (Thermofisher), targeting the bait-BirA, and streptavidin-DyLight 594 conjugate (Thermofisher), targeting the biotinylated proteins, were diluted at 1:300 and 1:1000 in blocking buffer, respectively and incubated with fixed cells for 30 minutes at room temperature. Cells were then washed, counterstained with DAPI, mounted on the slide using antifade vectashield mountant, and imaged using Leica SP8 Confocal with Lightning Deconvolution. Colocalization quantification was performed for the deconvolved images using Fiji’s (ImageJ 1.52p) Coloc_2 analysis tool between the 650 (anti-flag) and 594 (Streptavidin) channels (Schindelin, Rueden, Hiner, & Eliceiri, 2015). This tool generates a comprehensive report for evaluating pixel intensity colocalization of two channels by various methods such as Pearson’s Coefficient (range: - 1.0 to 1.0), Manders’ Colocalization Coefficients (MCC, range: 0 to 1.0), and Li’s Intensity Correlation Quotient (IQC, range: −0.5 to 0.5) (Li, 2004; Manders, Verbeek, & Aten, 1993). Background pixel intensity was subtracted using Fiji’s rolling ball algorithm and a region of interest (ROI). Thresholds were determined using Coloc_2’s bisection method, which is further used to adjust for background. Above threshold metrics were reported.

### RNAi knockdown experiment

An orthogonal siRNA approach called esiRNA (Sigma Aldrich) was employed to target PDIA4, PDIA6, and ERp44. HEK293 cell expressing SERPINC1-BirA and SERPINA1-BirA were seeded at 1X10^5^ cells/well in 12-well plates with complete medium and reverse transfected with 144 ng of the appropriate PDI specific esiRNA, Luciferase esiRNA as a negative control, or KIF11 esiRNA as positive control using Lipofectamine RNAiMAX (Invitrogen). All transfections were performed according to the manufacturer’s guidelines. Each experiment was done in triplicates and targeted gene knockdown by esiRNA was allowed to occur for 96 hrs. Culture supernatants and cell pellets were then harvested, clarified by low-speed centrifugation, and then aliquoted and stored at −80°C for further experiments.

### Western blotting

To validate the secretion of bait-BirA proteins, supernatants of cultures expressing fusion proteins were collected, centrifuged to remove cell debris, and 30 μl were loaded on SDS-PAGE gel for electrophoresis. The resolved proteins were then transblotted to nitrocellulose membranes using the Trans-Blot Turbo Transfer System from Bio-RAD. The membrane was blocked with 5% skim milk in TBST and probed with HRP-conjugated anti-flag mouse monoclonal antibody (Thermofisher) diluted at 1:10000 in the blocking buffer. The membrane was washed, and Clarity Western ECL Substrate was added. Proteins’ bonds were visualized using G:Box Gel Image Analysis Systems (SYNGENE). For staining of intracellular biotinylated proteins, cells were grown in complete medium supplemented with 50 μM biotin, lysed by RIPA buffer, and protein content was quantified using Bradford assay. 20 ug of total protein was loaded, resolved and transblotted as described earlier. The membrane was blocked by 3% BSA in TBST and probed with HRP-conjugated streptavidin diluted in blocking buffer at 1:2000 ratio. For visualizing the proteins’ bands, the same Clarity Western ECL Substrate was used.

Quantitative western blots were used to determine knockdown (KD) efficiency as well as impact on SERPIN secretion. For KD efficiency, cell pellets from each KD experiment were lysed with RIPA buffer, and approximately 25μg of protein lysate was loaded onto the SDS-PAGE gel for PDIA4, PDIA6, and ERp44 experiments. Resolved proteins were transblotted onto separate nitrocellulose membranes as described earlier. Each membrane was blocked with Intercept (TBS) Blocking Buffer from LI-COR and probed with Rabbit anti PDIA4 (1:2000), PDIA6 (1:2000), or ERp44 (1:2500) monoclonal antibodies, respectively. A housekeeping protein in each lysate was targeted for normalization with either a Mouse anti Alpha-tubulin (1:20,000) or anti Beta-actin (1:10,000) monoclonal antibodies. For visualization, IR imaging was utilized by staining with LI-COR Goat anti Mouse conjugated to 680 (1:15,000) and anti-Rabbit conjugated to 800 (1:15,000). Images were taken and analyzed with Image Studio Lite Version 5.2 for relative quantification. Impact on SERPIN secretion was determined from the aliquots of clarified supernatants from esiRNA transfected cultures using the same blotting method as above. 30 μl of supernatants were loaded on SDS-PAGE gel, and transblotted onto nitrocellulose membrane. Transblotted membrane was probed using Rabbit anti Flag (Proteintech, 1:800) followed by Goat anti Rabbit 800 (1:15,000). KD efficiency and the effect of PDI knockdown on secretion of the model proteins was measured in comparison with the negative control of each cell line transfected with Luciferase esiRNA as negative control (see above).

### Mass Spectrometry

Cells were grown in 245 mm plates (one plate per biological replicate in triplicate) to approximately 70% confluence in complete media and then incubated for 24 h with 50 μM biotin. Cells were harvested and washed twice in cold PBS, lysed with vigorous shaking (20 Hz) in 8M urea, 50mM ammonium bicarbonate lysis buffer, extracted proteins were centrifuged at 14,000 x g to remove cellular debris and quantified by BCA assay (Thermo Scientific) as per manufacturer recommendations. Affinity purification of biotinylated proteins was carried out in a Bravo AssayMap platform (Agilent) using AssayMap streptavidin cartridges (Agilent), and the bound proteins were subjected to on-cartridge digestion with mass spec grade Trypsin/Lys-C Rapid digestion enzyme (Promega, Madison, WI) at 70°C for 2h. Digested peptides were then desalted in the Bravo platform using AssayMap C18 cartridges and the organic solvent was removed in a SpeedVac concentrator prior to LC-MS/MS analysis. Dried peptides were reconstituted with 2% acetonitrile, 0.1% formic acid, and analyzed by LC-MS/MS using a Proxeon EASY nanoLC system (Thermo Fisher Scientific) coupled to a Q-Exactive Plus mass spectrometer (Thermo Fisher Scientific). Peptides were separated using an analytical C18 Acclaim PepMap column 0.075 x 500 mm, 2μm particles (Thermo Scientific) in a 93-min linear gradient of 2-28% solvent B (80% acetonitrile, 0.1% formic acid) at a flow rate of 300nL/min. The mass spectrometer was operated in positive data-dependent acquisition mode. MS1 spectra were measured with a resolution of 70,000, an AGC target of 1e6 and a mass range from 350 to 1700 m/z. Up to 12 MS2 spectra per duty cycle were triggered, fragmented by HCD, and acquired with a resolution of 17,500 and an AGC target of 5e4, an isolation window of 1.6 m/z and a normalized collision energy of 25. Dynamic exclusion was enabled with a duration of 20 sec.

### MS data Analysis

All mass spectra were analyzed with MaxQuant software (Tyanova, Temu, & Cox, 2016) version 1.5.5.1. MS/MS spectra were searched against the Homo sapiens Uniprot protein sequence database (version January 2018) and GPM cRAP sequences (commonly known protein contaminants). Precursor mass tolerance was set to 20ppm and 4.5ppm for the first search where initial mass recalibration was completed and for the main search, respectively. Product ions were searched with a mass tolerance 0.5 Da. The maximum precursor ion charge state used for searching was 7. Carbamidomethylation of cysteines was searched as a fixed modification, while oxidation of methionines and acetylation of protein N-terminal were searched as variable modifications. Enzyme was set to trypsin in a specific mode and a maximum of two missed cleavages was allowed for searching. The target-decoy-based false discovery rate (FDR) filter for spectrum and protein identification was set to 1%. Enrichment of proteins in streptavidin affinity purifications were calculated as the ratio of intensity. To remove the systematic biases introduced during various steps of sample processing and data generation, dataset were normalized using the LOESS method (Smyth, n.d.) integrated into Normalyzer (Chawade, Alexandersson, & Levander, 2014). Perseus software (Tyanova & Cox, 2018) was employed for data preparation, filtering, and computation of differential protein abundance. The DEP package (Zhang et al., 2018) was used to explore whether missing values in the dataset are biased to lower intense proteins. Left-censored imputation was performed using random draws from shifted distribution. A Student’s t-test with a multi-sample permutation-based correction for an FDR of 0.05 was employed to identify differentially expressed proteins using log2 transformed data.

### Detection of significant interactions

The threshold for significant interactions was determined using the known secretory pathway components as a gold standard. We set the cutoffs for FDR at 0.1 and removed all interactors with negative fold changes, as this optimizes the enrichment of known secretory pathway components among the significant interactors. The enrichment for two independent secretory pathway-related gene sets also peaked around the cutoffs set through the gene set of known secretory pathway components, suggesting the optimal cutoffs are robust to the gold standards chosen.

### Estimation of preferential interaction between protein features and interactors with a Bayesian modeling framework

To identify patterns of interactions between individual bait-BirA proteins and their interactors, we first obtained and summarized several protein features across model proteins. The protein properties considered include shared structural motifs (Blatch & Lässle, 1999), known sites of PTM from Uniprot and phosphosite (Blatch & Lässle, 1999; Hornbeck et al., 2015). The complete protein feature composition for each of the model proteins is illustrated in (Fig. S4). Given the interactions between the SecPs and their interactors, we can predict the important structural features implicated in the interactions between the SecPs and a given SecM. The interactions between the BirA-fused samples and the secretory pathway interactors can be pooled according to shared properties of the SecPs to reveal interdependencies between components of the secretory pathway and their products.

To test if some secretory machinery components preferentially interact with certain protein features, we first calculated the effective total frequency (*δ_f,g_*) of interactions between each feature-gene pair (*f,g*) by going through every SecP in our data and counting the number of times this feature occurs in a bait-BirA protein p (*f_p_*).

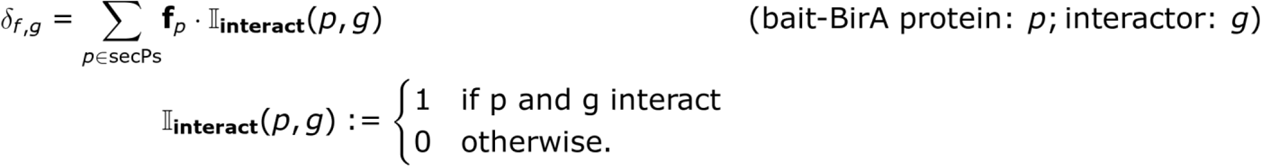

The number *f_p_* is added to the total frequency, *δ_f,g_*, only when *p* and *g* interact (when *II_interact_(p,g)* = 1). This effective interaction frequency, *δ_f,g_* essentially linked the features pooled across the model proteins to a given interactor *g*, taking us closer to estimating interaction affinity between a feature and an interactor. To further account for SecP and feature promiscuity, we implement an estimate of the tendency for f and g to interact. The interaction synergy *s_f,g_*, the tendency for an interactor *g* to interact with a feature *f* more than expected by chance, was estimated by a Bayesian modeling approach. More specifically, we seek to decouple the interaction synergy from the observed interaction frequency *δ_f,g_* by modeling the interaction frequency *δ_f,g_* with a Poisson regression.

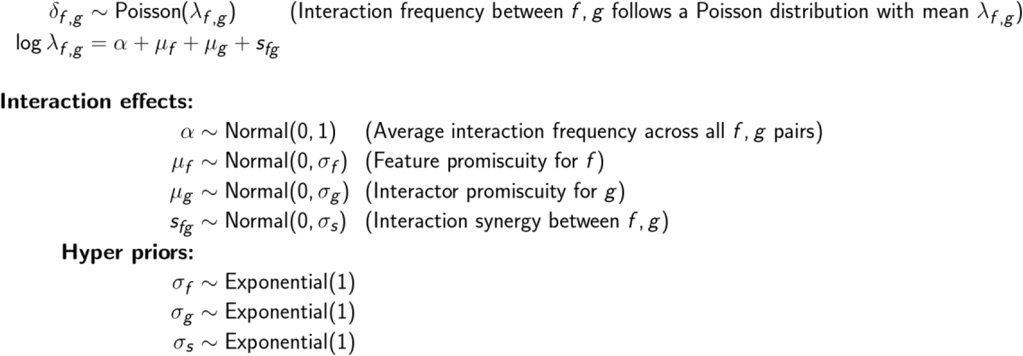

The mean for the Poisson distribution is parameterized by the sum of feature promiscuity (number of SecMs *g* connected to a feature) *μ_f_*, interactor promiscuity (number of SecPs with a feature *f* interacting with a SecM) *μ_g_*, the interaction synergy *s_f,g_* and an intercept variable. The feature promiscuity *μ_f_* quantifies the probability of a feature *f* to partake in an interaction with any secMs. Likewise, the interactor promiscuity *μ_g_* measures the tendency for a secM *g* to interact with any features. After subtracting out the *μ_f_* and *μ_g_* from the log-transformed mean *λ_f,g_* for the Poisson distribution, we arrive at the coefficient of interest-- the interaction synergy *s_f,g_* between *f* and *g*. It quantifies the degree to which *f* and *g* interact more than by random chance. In previous work, this approach has correctly estimated epistasis intensity(Shen et al., 2017). To better regularize the promiscuities, their Bayesian priors are all normally distributed around 0, with their variances parameterized by the hyper priors *σ_f_, σ_g_* and *σ_s_* which follow an exponential distribution. The intercept *α* is parameterized by a standard normal distribution. We used the *rethinking* R package (McElreath, 2020) to construct the model and sample the coefficients.

## Results

### BioID can successfully tag proteins colocalized with secreted proteins

We first investigated if intracellular PPIs between each SecP and their supporting SecMs can be measured using the BioID method. To do this, each bait-BirA was expressed in HEK293 cells using the Flp-In™ system (see materials and methods) for targeted integration of the transgenes into the same genomic locus to ensure comparable transcription rates of each transgene. Variations in mRNA level caused by random integration can trigger adaptive response such as the unfolded protein response in some cell lines which reciprocally alters the active PPIs network involved in the secretion. We observed successful secretion of bait-BirA proteins into culture supernatant, evaluated by Western blot (Fig. 2a). Thus, the BirA fusion did not prevent secretion of the model proteins, and it is expected that they enter the secretory pathway where they are processed and packaged for secretion. We also verified the biotinylation profile by western blot for each cell line in the presence and absence of biotin. The biotinylation profile of the bait-BirA cells is different when biotin is added to the culture with substantial increased biotinylation of specific proteins, while no obvious change is observed for WT (Fig. 2b), suggesting that BioID2 successfully tagged specific proteins within the cells.

**Figure 2.**
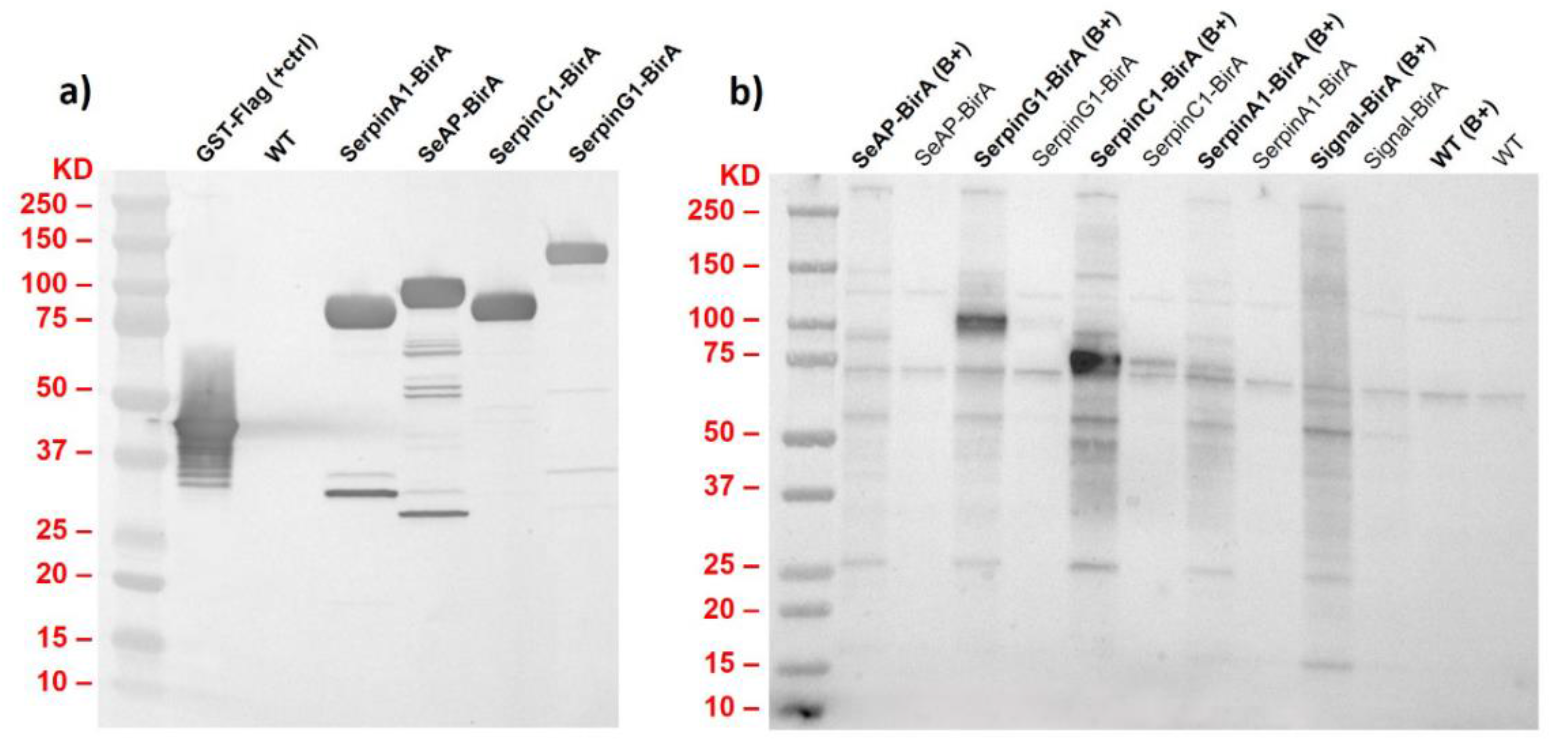
Expression of bait-BirA proteins results in a substantial increase in biotinylated proteins. a) Successful secretion of the bait-BirA proteins into the culture supernatant was evaluated by Western blot using HRP-anti-flag antibody. b) The immunoblotting biotinylation profiling of the model proteins and WT control in HEK293 cells with HRP-streptavidin. When the BirA domain was fused to the model proteins, biotin addition led to the biotinylation of a subset of proteins (B+) which are not seen in WT or absence of biotin. This demonstrates that the BioID labeling system tags interactions as secreted proteins are synthesized and trafficked through the secretory pathway. A few endogenously biotinylated proteins appear in the absence of biotin and in the WT.

Colocalization of the bait-BirA proteins and the biotinylated proteins was then studied by multicolor co-immunofluorescence microscopy to test whether biotinylated proteins are actual partners of the model proteins. The results demonstrated successful labeling of the interactors by BirA through colocalization of the biotin-labeled proteins and bait-BirA, while WT did not show increased biotinylation under the same experimental conditions (Fig. 3). To quantify the colocalizations, we calculated different colocalization metrics (see methods) from the images and compared to the WT (Table 1 and Fig. S1), and the results confirmed the specificity of the BioID labeling system to tag the proximal proteins (Li’s ICQ value closer to 0.5 and Pearson’s R value and Manders’ Colocalization Coefficients closer to 1 demonstrate a dependent protein staining pattern between the red and green channels).

**Table 1.** Colocalization metrics between the 650 (anti-flag) and 594 (Streptavidin) channels in Fig. S1. There is little correlation reported for the WT, whereas there is much higher correlation for clones containing BirA.

| Cells | Pearson's R value | Li's ICQ value | Manders' Coeff. (tM<sub>g</sub>,tM<sub>r</sub>) |
| --- | --- | --- | --- |
| WT | 0.22 | 0.219 | 0.812, 0.518 |
| SERPINA1-BirA | 0.74 | 0.318 | 0.962, 0.888 |
| SERPING1-BirA | 0.73 | 0.260 | 0.999, 0.914 |
| SERPINC1-BirA | 0.88 | 0.392 | 0.991, 0.982 |
| SeAP-BirA | 0.93 | 0.392 | 0.952, 0.935 |
| Signal-BirA | 0.71 | 0.308 | 0.841, 0.791 |

**Figure 3.**
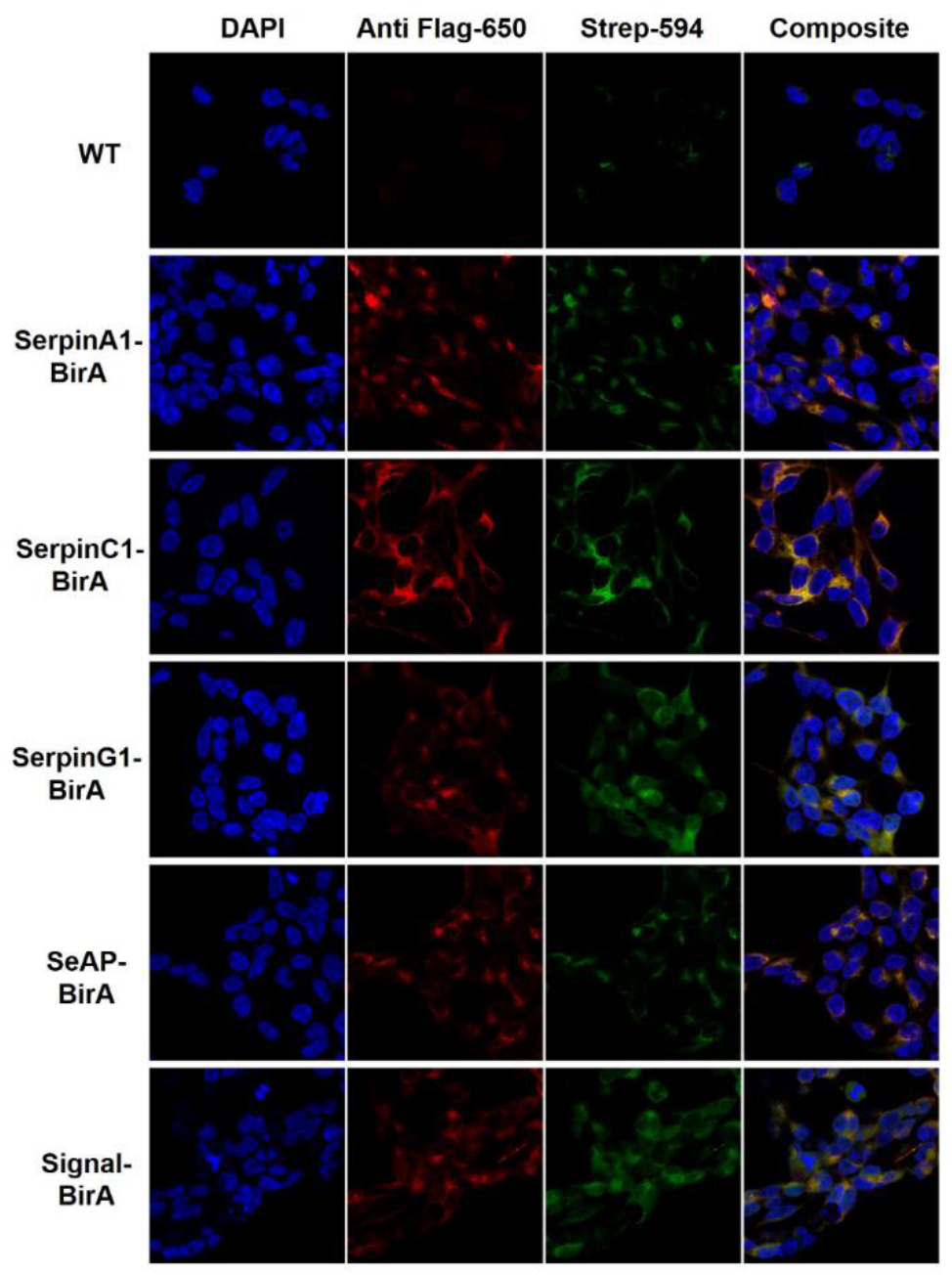
Bait-BirA fusion proteins are colocalized with biotin-staining. Co-Immunofluorescence demonstrated the intracellular colocalization of the biotin-labeled proteins (stained with Streptavidin-Dylight 594 and illustrated in green color) and bait-BirA (stained with anti-flag monoclonal antibody-Dylight 650 and illustrated in red color), while WT did not show increased biotinylation under the same experimental conditions.

### WT cells revealed endogenous biotinylation landscape

After successful tagging of the proximal proteins we aimed to identify the interactions with each bait protein. For this, cells were lysed, and biotinylated proteins were purified using streptavidin. Purified proteins were digested by trypsin, and peptides were subjected to LC-MS/MS, and the biotinylated proteins in samples were mapped using MaxQuant (Tyanova et al., 2016). Differentially biotinylated proteins were then identified in each sample compared to the WT using Perseus (Tyanova & Cox, 2018). When implemented in a WT control cell line, we identified proteins that are biotinylated endogenously along with bona fide interactors. These include proteins that bind biotin as a cofactor such as carboxylases which is problematic with streptavidin-based protein detection (Tytgat et al., 2015). Although the extent to which the general endogenous biotinylation has not been systematically quantified, the biotinylated proteins isolated from the WT sample showed considerable overlap with interacting proteins detected in other model protein samples, suggesting endogenous biotinylation may be more pervasive than previously believed. Among the top differentially biotinylated proteins in bait-BirA samples, the bait proteins showed the highest log-fold change (LFC) (Fig. 4 and Table S1). This observation is expected because the bait protein is a potential substrate for BirA located in the closest vicinity of the enzyme and is considered as evidence to show the biotinylation system is working properly in the cell.

**Figure 4.**
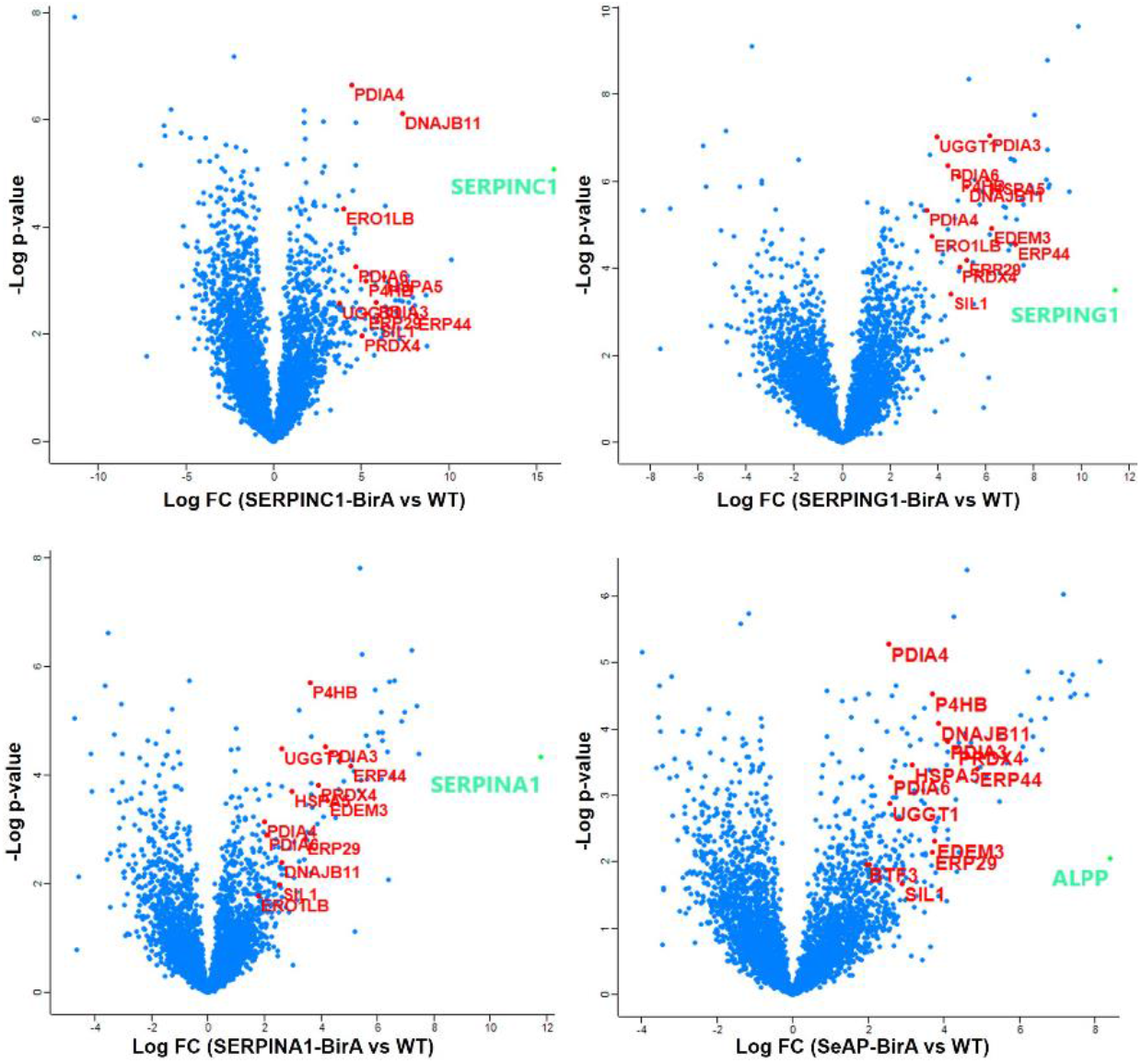
Dozens of proteins show significantly increased biotinylation after expression of bait-BirA proteins. Volcano plot showing the distribution of the quantified biotinylated proteins by MS according to p-value and fold change. As depicted the bait-protein significantly showed the highest fold change compared to WT almost in all cases, indicating the capability of the BioID labeling system to tag the in vivo interactions within the live cells. The key interactors involved in disulfide bond and N-glycosylation formation (see text) are highlighted in red. SeAP is a truncated form of Alkaline phosphatase, placental type (ALPP).

### Interactors are enriched for secretory pathway components and co-secreted proteins

In theory, the model proteins would come in frequent contact with members of the secretory pathway, and other co-secreted proteins. To determine if our setup captures the secretory pathway-related proteins more than by random chance, we analyzed the enrichment for 3 independent secretory pathway-relevant gene sets at various fold change and p-value thresholds. We saw a significant enrichment for the secretory pathway machinery, secretory-resident and co-secreted proteins among probable PPIs across all model protein samples (Fig. 5a and Fig. S3), with peak enrichment occurring at the top 100-300 most significant interactors by positive fold change. This corresponds to a significance cutoff of a fold change of 3 or greater enrichment and an adjusted p-value < 0.1, in model proteins compared to WT control (Fig. S2 for all thresholds and Fig. 5b for all significant interactions) for each secreted protein (see methods). The secretory machinery components are more enriched among the top 300 hits for all model proteins than other co-secreted proteins, suggesting more frequent interactions between the secretory pathway machinery and their products than the crosstalk between co-secreted proteins. Probable PPIs detected in all model proteins (n=19) and hits shared among all SERPIN gene products are significantly enriched for proteins involved in protein folding (Fig. 5b). Indeed, molecular chaperones are highly promiscuous when assisting protein folding due to their inherent flexibility (Mayer, 2010). Apart from the shared interactions, PPIs for each model protein differ substantially. Thus, the question remained if these private interactions correspond to unique properties of each model protein.

**Figure 5.**
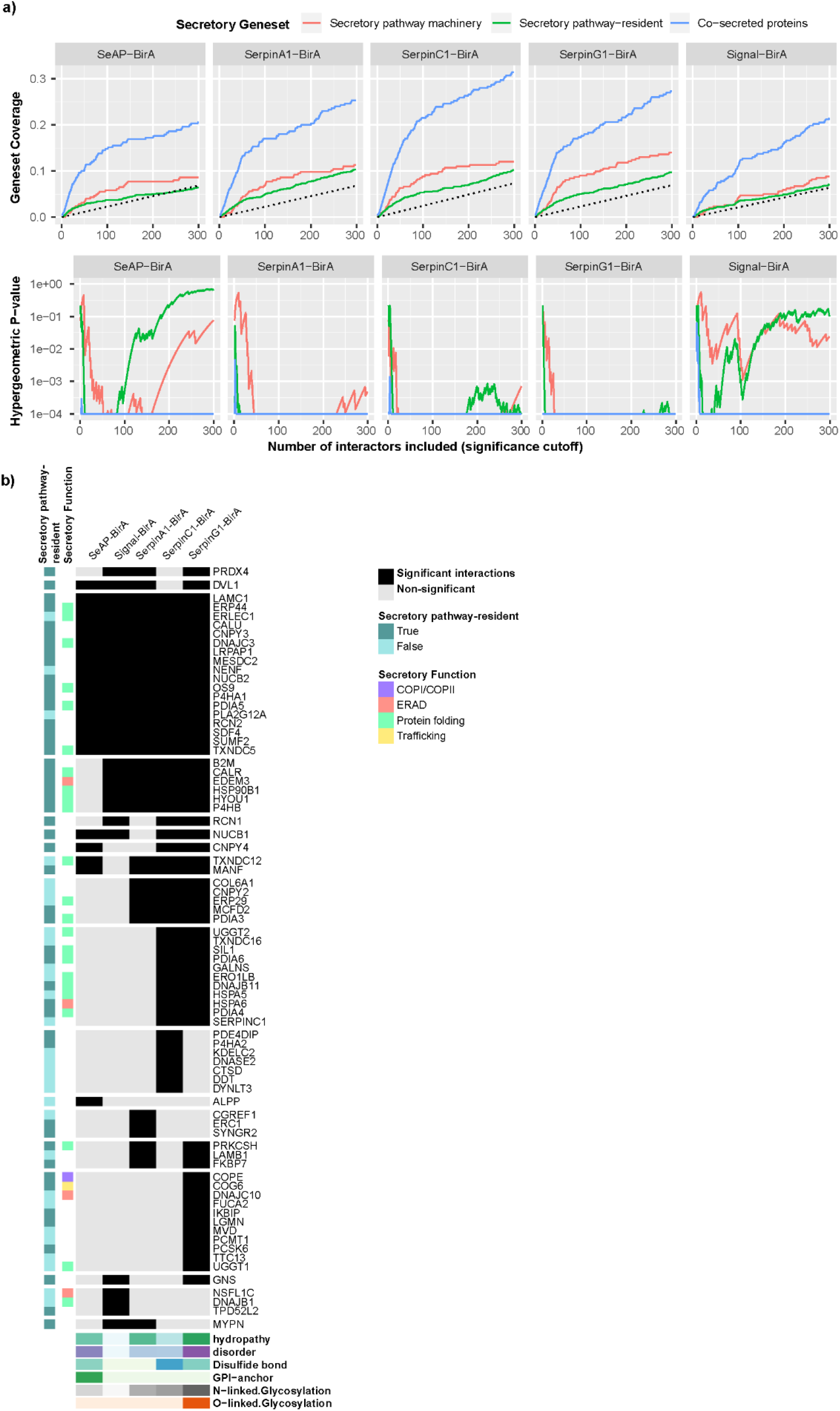
Interacting proteins are enriched for secretory pathway machinery. (a) To determine if significant interactions enrich for secretory pathway-related genes, we performed an iterative enrichment analysis in which we included the interactors with the greatest fold changes first and iteratively added interactors with lower fold changes. The y-axis indicates the overall coverage of 3 secretory pathway-related gene sets and the x-axis the significance cutoffs (rank ordered by fold change). The coverage of the gene set (top) along with their corresponding hypergeometric enrichment p-value (bottom) are shown, denoting the probability of obtaining an overlap larger than the one observed if no enrichment existed. The top 300 hits for each secretable BirA sample (Fig. S3 for all hits) showed significant enrichment of the secretory pathway components and co-secreted proteins among the most significant hits for all samples except Signal-BirA (which is a lone secreted BirA and not a mammalian secreted protein). (b) Quantified interactions between interactors (y-axis) and the model proteins (x-axis), where the shadowed entries indicate significant interactions. The features of the model proteins, detailed in Fig. S4, are summarized for each model protein on the bottom panel and the secretory pathway attributes for the interactors are labeled on the left.

### Private interactors reflect post-translational and structural features of model proteins

PPIs in the secretory pathway mediate the folding, modification and transportation of secreted proteins (Bonifacino & Glick, 2004; Calakos, Bennett, Peterson, & Scheller, 1994; Ikawa et al., 1997; Pearl & Prodromou, 2006; Watanabe et al., 2019). Incidentally, co-expression analysis has linked certain PTMs across the secretome to the expression of their responsible enzymes. For example, PDIs are consistently upregulated in tissues secreting disulfide-rich proteins (Feizi et al., 2017). As the bait-BirA proteins differ in structural composition and PTMs (Fig. S4), we wondered if bait-BirA proteins with shared features have higher affinity for specific interactors. More specifically, we hypothesize that proteins requiring a specific PTM would preferentially interact with the secretory machinery components responsible for the PTM synthesis. To test if such preferential interaction exists, we summarized various PTM and structural properties across model proteins and analyzed their associations with the corresponding secretory machinery using a Bayesian modeling framework (see methods). Among the studied PTMs, bait-BirA proteins with disulfide bonds and N-linked glycans demonstrated higher affinity towards specific interactions (Fig. 6a) that are known to help secretion of proteins with the corresponding PTMs. Thus, we analyzed the detected interactions associated with glycosylation, disulfide bond addition, and protein folding.

**Figure 6.**
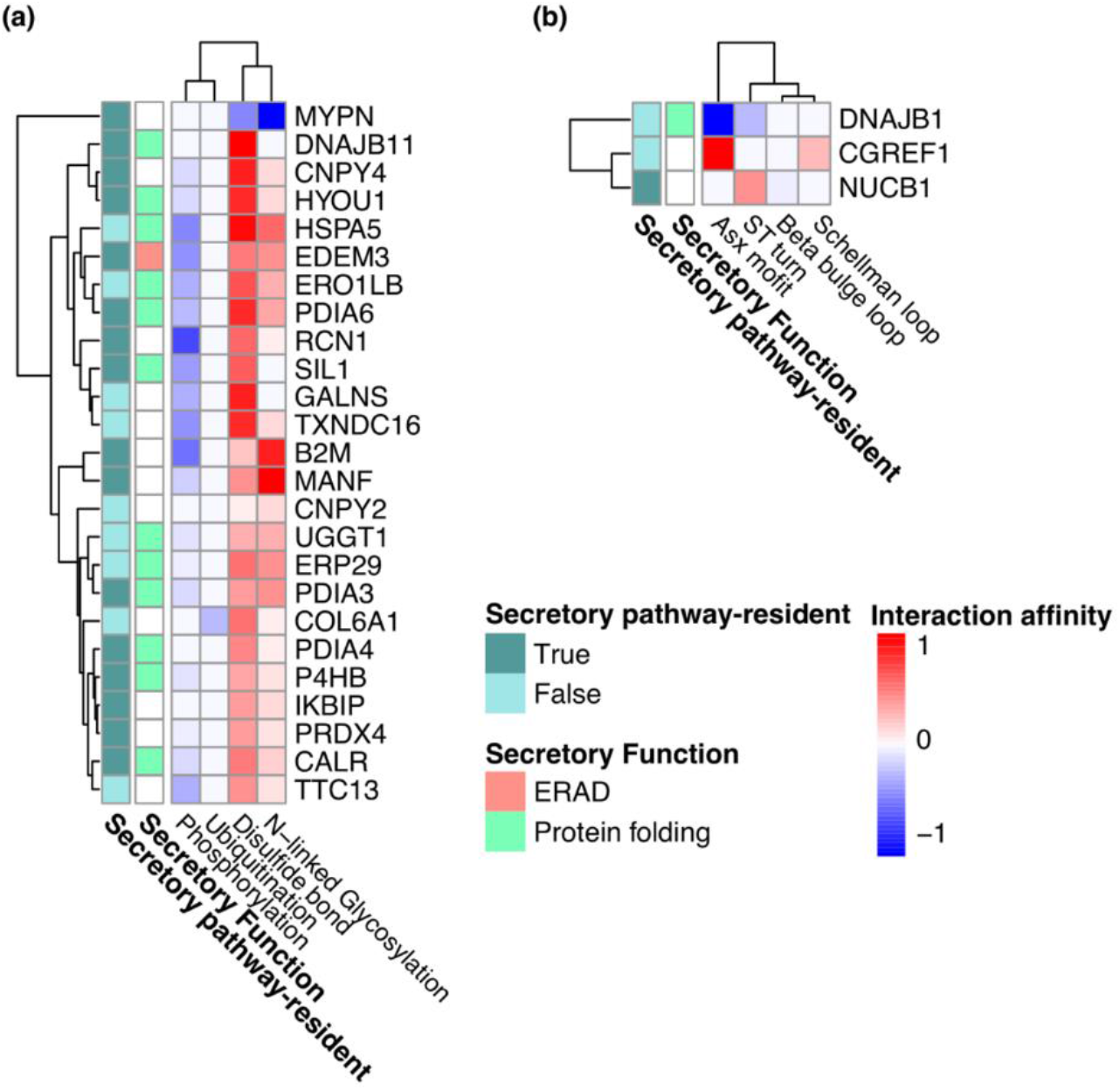
Detected interactors correlate with protein features. Interactors were associated with specific (a) PTMs and (b) structural features of model proteins. The heatmap shows the standardized interaction affinities estimated between certain interactors and PTMs or structural features across all model proteins (see methods). Only interactors having significant associations with model protein features are shown.

### Proteins with increased glycosylation are associated with quality control pathways

We detected significant interactions in the Calnexin/Calreticulin cycle and related processes for more heavily glycosylated proteins (Fig. 6a). For example, the glycosylated baits interacted with calreticulin (CALR), a calcium-binding chaperone that promotes folding, oligomeric assembly, and quality control of glycoproteins in the ER(Nauseef, McCormick, & Clark, 1995). They also interacted with UGGT1, which recognizes glycoproteins with minor folding defects and reglucosylates single N-glycans near the misfolded part of the protein. Reglucosylated proteins are then recognized by CALR for recycling to the ER and refolding or degradation (Ferris, Jaber, Molinari, Arvan, & Kaufman, 2013). Two members of the PDI family, PDIA3 and ERp29, which form a complex with calreticulin/calnexin, also showed association with N-glycosylated baits (Fig. 6a) suggesting their role in glycoprotein folding and quality control. Calnexin/Calreticulin-PDIA3 complexes promote the oxidative folding of nascent polypeptides (Sakono, Seko, Takeda, & Ito, 2014) and ERp29 promotes isomerization of peptidyl-prolyl bonds to attain the native polypeptide structure (Sakono et al., 2014; Tannous, Pisoni, Hebert, & Molinari, 2015). Proteins with chaperone activity, such as HSPA5 (Fig. 6a), were also found to interact with N-linked glycan-containing bait-BirA. HSPA5 is a component of the glycoprotein quality-control (GQC) system which recognizes glycoproteins with amino acid substitutions, and targets them for ER-associated degradation (ERAD) (Ferris, Kodali, & Kaufman, 2014). EDEM3, another interactor associated with the N-glycan containing proteins (Fig. 6a), is a glycosyltransferase involved in ERAD mediated degradation of glycoproteins by catalyzing mannose trimming from Man8GlcNAc2 to Man7GlcNAc2 in N-glycans (Ninagawa et al., 2014). Given that most of these molecular chaperones and enzymes are involved in ERAD-mediated degradation of the misfolded glycoproteins, these findings suggest the quality control pathways are critical for synthesizing and secreting proteins with N-linked glycans.

### Disulfide bond formation is rate-limiting in protein secretion

Several members of the PDI family including P4HB, PDIA3, PDIA4 and PDIA6 significantly interacted with model-BirAs containing more disulfide bonds (Fig. 6a). These enzymes catalyze the formation, breakage and rearrangement of disulfide bonds through the thioredoxin-like domains (Kozlov, Määttänen, Thomas, & Gehring, 2010). The identification of various PDIs highlights the importance of the oxidative folding enzymes in protein folding and maintaining stability that can limit the efficiency of protein secretion. The proteins with more disulfide bonds also interact with major ER chaperones HSPA5 and DNAJB11, a co-chaperone of HSPA5, that play a key role in protein folding and quality control in the ER lumen (Ng, Watowich, & Lamb, 1992; Yu, Haslam, & Haslam, 2000), highlighting their important role in secretion of the disulfide bond enriched proteins. The PDI, ERp44 showed the highest association (LFC > 8, Fig. S5) with disulfide bond-enriched proteins i.e. SERPINC1 and SERPING1 although it also showed considerable interaction with SERPINA1-BirA and a less strong interaction with SeAP-BirA. ERp44 mediates the ER retention of the oxidoreductase Ero1α (an oxidoreductin that reoxidizes P4HB to enable additional rounds of disulfide formation) through the formation of reversible mixed disulfides (Anelli et al., 2003). Hence, the association of ERp44 and disulfide bonds highlights the importance of the thiol-mediated ER protein retention in disulfide bond formation, particularly when the secretory pathway is loaded with proteins with more disulfide bonds. In addition, ERO1LB, PRDX4 and SIL1 were ER-localized enzymes that were associated with disulfide bond formation. ERO1LB efficiently reoxidizes P4HB (Mezghrani et al., 2001), PRDX4 couples hydrogen peroxide catabolism with oxidative protein folding by reducing hydrogen peroxide (Zito, 2013), and SIL1 can reverse HSPA5 cysteine oxidation which alters its chaperone activity to cope with suboptimal folding conditions (Siegenthaler, Pareja, Wang, & Sevier, 2017). The identification of these oxidoreductase enzymes highlights the importance of ER redox homeostasis in disulfide bond formation and protecting cells from the consequences of misfolded proteins.

To validate the importance of specific PDI interactions that showed highest fold change and specificity of effect on productivity of proteins with more disulfide bonds, we knocked down PDIA4, PDIA6, or ERp44 in cells expressing either SERPINC1-BirA or SERPINA1-BirA, using an orthogonal RNAi approach, i.e. esiRNAs (Kittler, Heninger, Franke, Habermann, & Buchholz, 2005). HEK293 cells expressing SERPINC1-BirA or SERPINA1-BirA were transfected with esiRNA against PDIA4, PDIA6, ERp44, or LUC and KIF11 as negative/positive controls, respectively. Knockdown experiments successfully resulted in >70% reduction in cellular expression of the targeted PDIs (Fig. 7a). We observed a 42% reduction in SERPINC1-BirA secretion following knockdown of PDIA4 (p-value = 0.032), meanwhile a significantly lesser reduction of SERPINA1-BirA secretion was seen (Fig. 7b). If differences in KD efficacy between experiments are accounted for, normalization of SERPINs secretion by KD efficiencies resulted in 54% reduction of SERPINC1-BirA secretion following knockdown of PDIA4 (p-value = 0.026) while no significant reduction was seen in SERPINA1-BirA secretion (Fig. 7c). Neither knockdown of PDIA6 nor ERp44 produced a significant reduction in either SERPINC1-BirA nor SERPINA1-BirA secretion (Fig. 7b & 7c). BioID analysis indicated that PDIA4 interacts significantly with SERPINC1-BirA but not with SERPINA1-BirA, while PDIA6 did not show considerable interaction with SERPINA1 nor SERPINC1 (Fig. S5). Therefore, PDIA4 may work as a private interactor for SERPINC1 secretion as recognized by knockdown results. PDIA6 could potentially work as a private interactor for SERPING1 secretion as predicted by BioID (Fig. S5). ERp44 had a higher quantified fold change interaction with SERPINC1-BirA, but it also showed significant interaction with SERPINA1-BirA, SERPING1-BirA and less strongly with SeAP-BirA, suggesting ERp44 may function in a more public manner.

**Figure 7.**
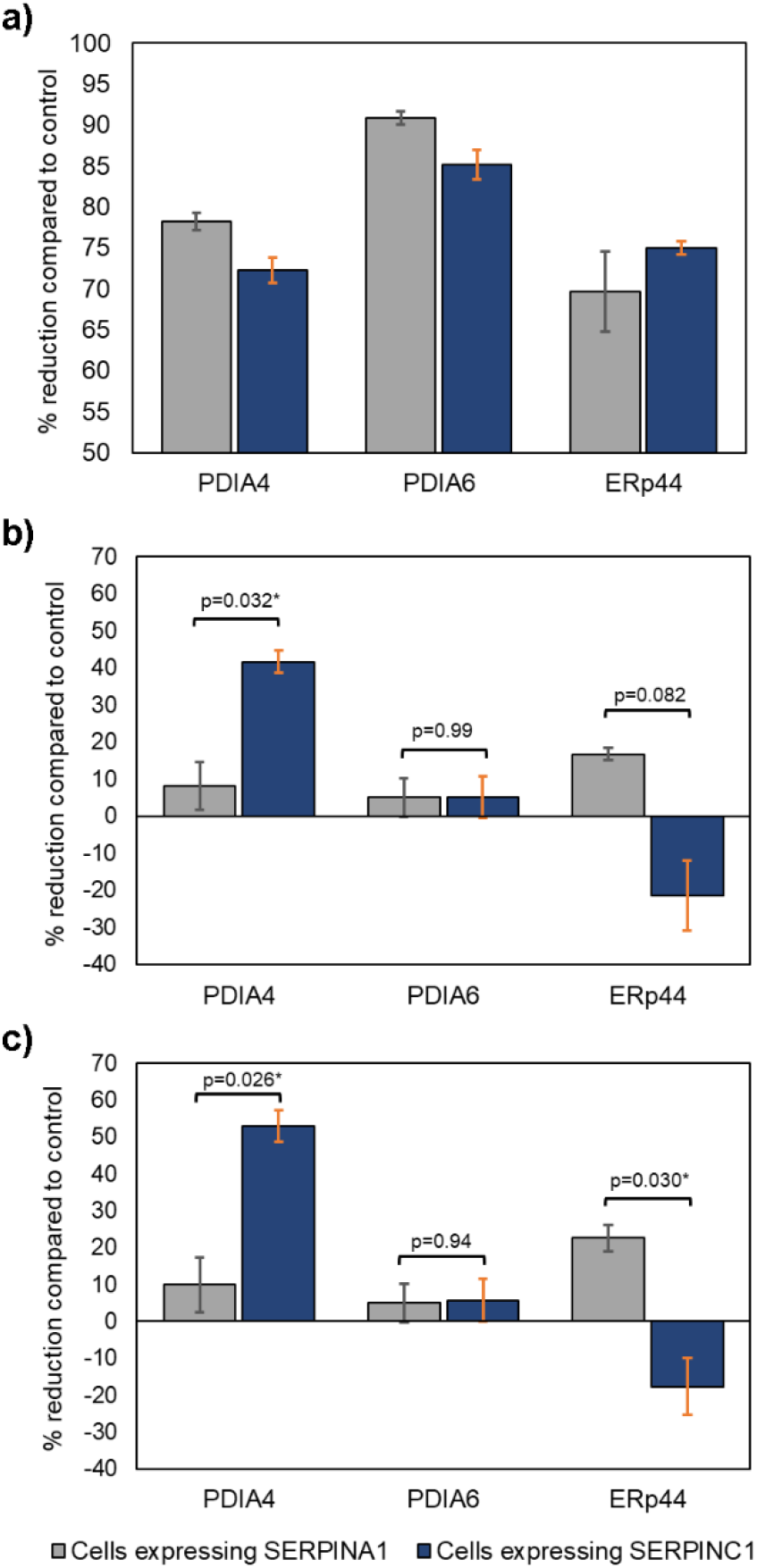
Effects of esiRNA mediated knockdown of isomerases PDIA4, PDIA6, and ERp44 on SERPIN secretion. Quantitative western blots were used to find the relative abundance of each PDI and SERPIN after esiRNA transfection in both cells expressing SERPINA1 and SERPINC1. We have reported these results as (a) KD efficiency of each PDI measured by percent reduction of PDI abundance compared to the negative control, (b) the effect of each PDI KD on SERPINA1 and SERPINC1 secretion, and (c) the normalized effects of PDI KD on SERPIN secretion using the KD efficiency, assuming greater KD results would result in a greater loss of SERPIN secretion. SERPINC1, which has a multitude of disulfide PTMs, shows a greater reduction in SERPIN production in PDIA4 KD experiments compared to SERPINA1, which does not have as many disulfide bridges in its structure. This is consistent with expected interactions found with BioID analysis.

### Identified PPIs are associated with structural motifs on bait proteins

In addition to PTMs, the Bayesian modeling framework found associations between SecP structural features and the SecMs (Fig. 6b). For example, model proteins depleted in the asx motif (Golovin & Henrick, 2008) showed higher tendency to interact with DNAJB1, a cytosolic molecular chaperone of the HSP40 family, possibly through retrotranslocation of the misfolded model proteins in ERAD. The asx motif impacts N-glycan occupancy of Asn-X-Thr/Ser sites, depending on the ability of the peptide to adopt an Asx-turn motif (Imperiali, Shannon, & Rickert, 1992; Imperiali, Shannon, Unno, & Rickert, 1992). As another example, NUCB1, a chaperone-like amyloid binding protein that prevents protein aggregation (Bonito-Oliva, Barbash, Sakmar, & Graham, 2017), interacted more strongly with our proteins with more ST-turns (Fig. 6b). ST turns occur frequently at the N-termini of α-helices (Doig, Stapley, Macarthur, & Thornton, 2008) and are regarded as helix capping features which stabilize α-helices in proteins (Aurora & Rosee, 1998). Thus, the enriched interaction of the NUCB1 with St-turn suggests that it can help stabilize folding of protein with a predominant α-helical secondary structure.

## Discussion

BioID has been used to profile the proteome of different cellular compartments and molecular complex systems (Varnaitė & MacNeill, 2016). However, this is the first time that BioID has been used to identify the proteome of the secretory pathway during recombinant protein expression. Numerous PPIs guiding the folding, modifications, and trafficking of the secreted and membrane proteins through the secretory pathway are transient (Nyfeler, Michnick, & Hauri, 2005; Schreiber, Haran, & Zhou, 2009), and therefore, cannot be captured by conventional methods such as co-immunoprecipitation. Consistent with previous studies (Sears, May, & Roux, 2019), these results showed the BioID can detect weak and transient interactions *in situ*, and therefore it is a powerful approach to study luminal processes involved in protein secretion. We found that disulfide bridge formation enzymes showed the strongest association with bait proteins enriched in disulfide bonds, supporting their critical roles in protein secretion and maintaining ER stability. A previous study on difficult to express (DTE) monoclonal antibodies showed less recognition by PDI impairs disulfide bridge formation within the antibody light chain (LC) which can initiate the intracellular degradation by the ubiquitin proteasome system via ERAD (Mathias et al., 2020). Thus, insufficient interaction between the secreted proteins with enriched disulfide bonds and PDIs can limit secretion efficiency and serve as a rate-limiting step for protein production. In another study, the tissue specific analysis of SecMs expression showed a positive correlation between the expression of P4HB and PDIA4 and liver tissue where numerous disulfide bond enriched proteins are secreted by hepatocytes (Feizi et al., 2017). These observations are clear evidence that suggests the tissue-specific fine-tuning of the PDI family expression level in response to the enrichment of the disulfide sites. Together, these results showed PDIs are actively involved in adaptive responses and secretion of proteins with dominant disulfide bonds that are crucial for restoring ER stability, and therefore, yielding the recombinant proteins. Given the associations between the SecMs and the features of the model proteins, we also hypothesized that SecMs preferentially interacting with bait-BirA proteins that carry certain structural features may be essential for the secretion of those proteins. While evidence linking SecMs to the structural motifs is lacking, many molecular chaperones selectively interact with certain sequence and structural elements to favor the particular folding pathways (Gidalevitz, Stevens, & Argon, 2013). For example, chaperones of the HSP70 family evolved to bind extended β strand peptides; interestingly, the associations identified between chaperones and asx motif and ST turn represent a novel association for further study.

While we show BioID works well for studying the synthesis of secreted proteins, we acknowledge that biotin-based methods have some limitations as well. Biotin is actively imported into the cytoplasm of cells and can freely diffuse to the nucleus, but it may not be as accessible in the secretory pathway, thus reducing labeling efficacy in that compartment (Kim & Roux, 2016). Here, we showed this challenge is not an insurmountable issue, in that the BioID2 construct successfully identified many expected luminal interactions. BioID2 requires less biotin supplementation and exhibits enhanced labeling of proximate protein (Kim et al., 2016), allowing for BioID to be introduced to new systems where biotinylation supplementation may not be easily accomplished (Sears et al., 2019). More recently, two promiscuous mutants of biotin ligase, TurboID and miniTurbo, have been developed to catalyze proximity labeling even with much greater efficiency (Branon et al., 2020) and therefore, can be considered as an effective method when proximal labeling of the endomembrane organelles is desired. It is also possible that the coverage of biotin labeling may be limited in the early processes in the secretory pathway due to the maturation process of the biotin ligase, i.e. biotinylation only can occur once the biotin ligase has been fully functional. Our results did reveal that there was coverage of the early secretory pathway interactions, but the possibility of missing interactions cannot be ruled out. So, BioID should be considered as a complementary method to other PPIs and systems biology approaches for complete characterization of endomembrane interactions.

In summary, we demonstrate here an approach to identify the protein interactions that synthesize and support secreted proteins, and thus define the product-specific secretory pathway. The identification of such machinery opens avenues for mammalian synthetic biology, wherein biotherapeutic production hosts can be rationally engineered to improve the titer and quality of diverse proteins in a client specific manner.

## Acknowledgments

This work was supported by generous funding from the Novo Nordisk Foundation provided to the Center for Biosustainability at the Technical University of Denmark (NNF10CC1016517, BGV) and from NIGMS (R35 GM119850, NEL). Light microscopy was performed at the UCSD School of Medicine Microscopy Core, which is supported by an NINDS P30 grant (NS047101). The authors would also like to thank Tune Wulff for support on pilot assays.

