## Supplementary figures for "In situ detection of protein interactions for recombinant therapeutic enzymes": Supplementary Materials.pdf

9500 Gilman Dr. MC 0760, Building BRF2, Room 4a16, La Jolla, CA 92093

**Short title:** In situ detection of protein interactions.

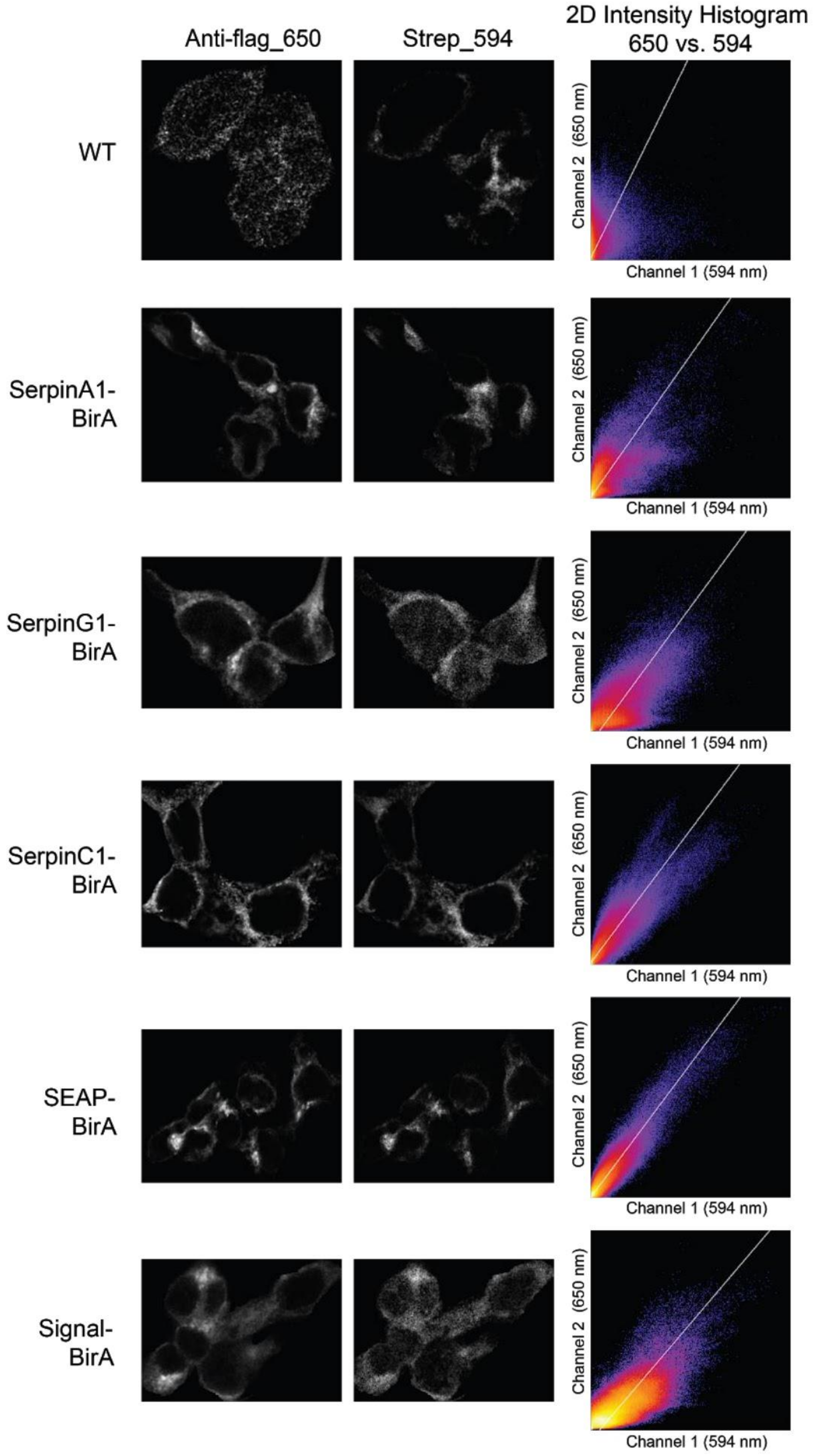


**Figure S1. Colocalization quantification between the 650 (Anti-flag) and 594 (Streptavidin) channels.** Max projections of anti-flag_650 and streptavidin_594 fluorescent channels with corresponding intensity histogram for one of various regions tested for each clone. These intensity plots were created using the same ImageJ Coloc2 tools as the colocalization metrics in Table 1. The relative linearity of the intensity graph demonstrates colocalization for the clones expressing the birA fusions and biotin labeling for each recombinant clone compared to the WT.

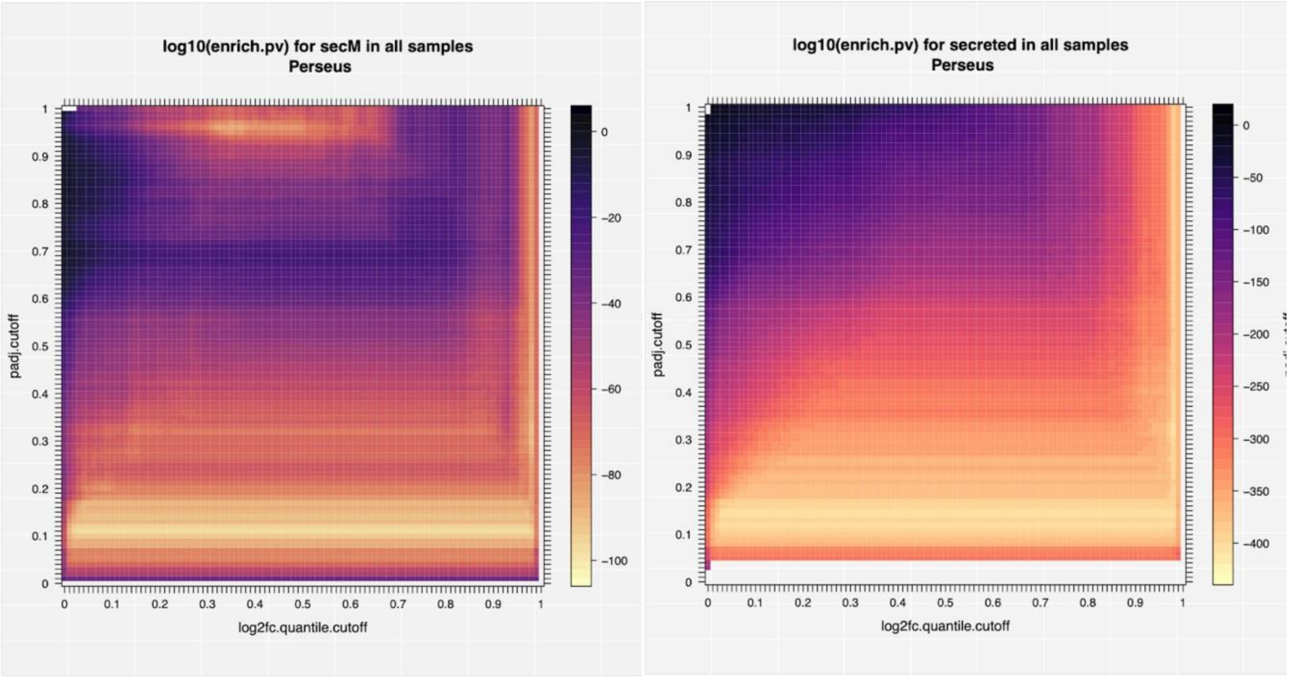


**Figure S2. As Perseus reported both log fold changes and p-values, we can optionally threshold significant interactors based on both criteria.** By extending Fig. 5 to include cutoffs for p-values, the enrichment ROC curves became a 2D heatmaps, where the x-axis, y-axis and color represent the fold change cutoffs, p-value cutoffs and the enrichment hypergeomic p-values respectively. As demonstrated by the near vertical and horizontal patterns near significant p-values and log fold changes (top <10%), these two criteria are mutually redundant for significant interactors.

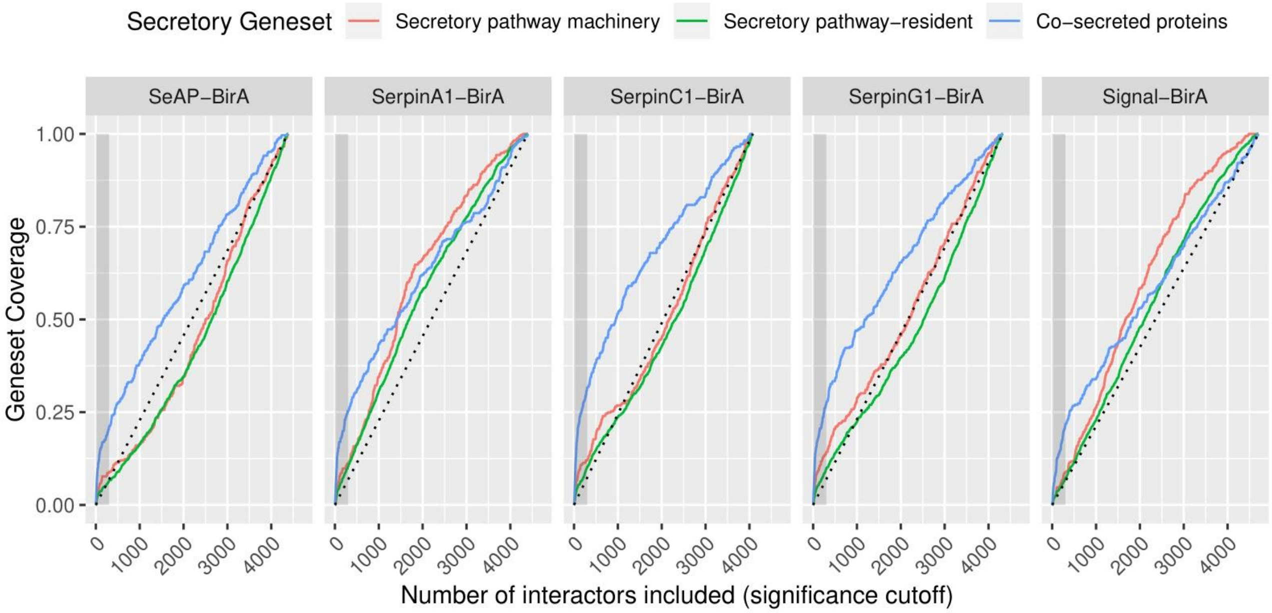


**Figure S3. Iterative enrichment of 3 secretory-related gene sets of all hits for each bait-BirA sample**. The shadowed region in each sample indicates the top hits zoomed-in in Fig. 5.

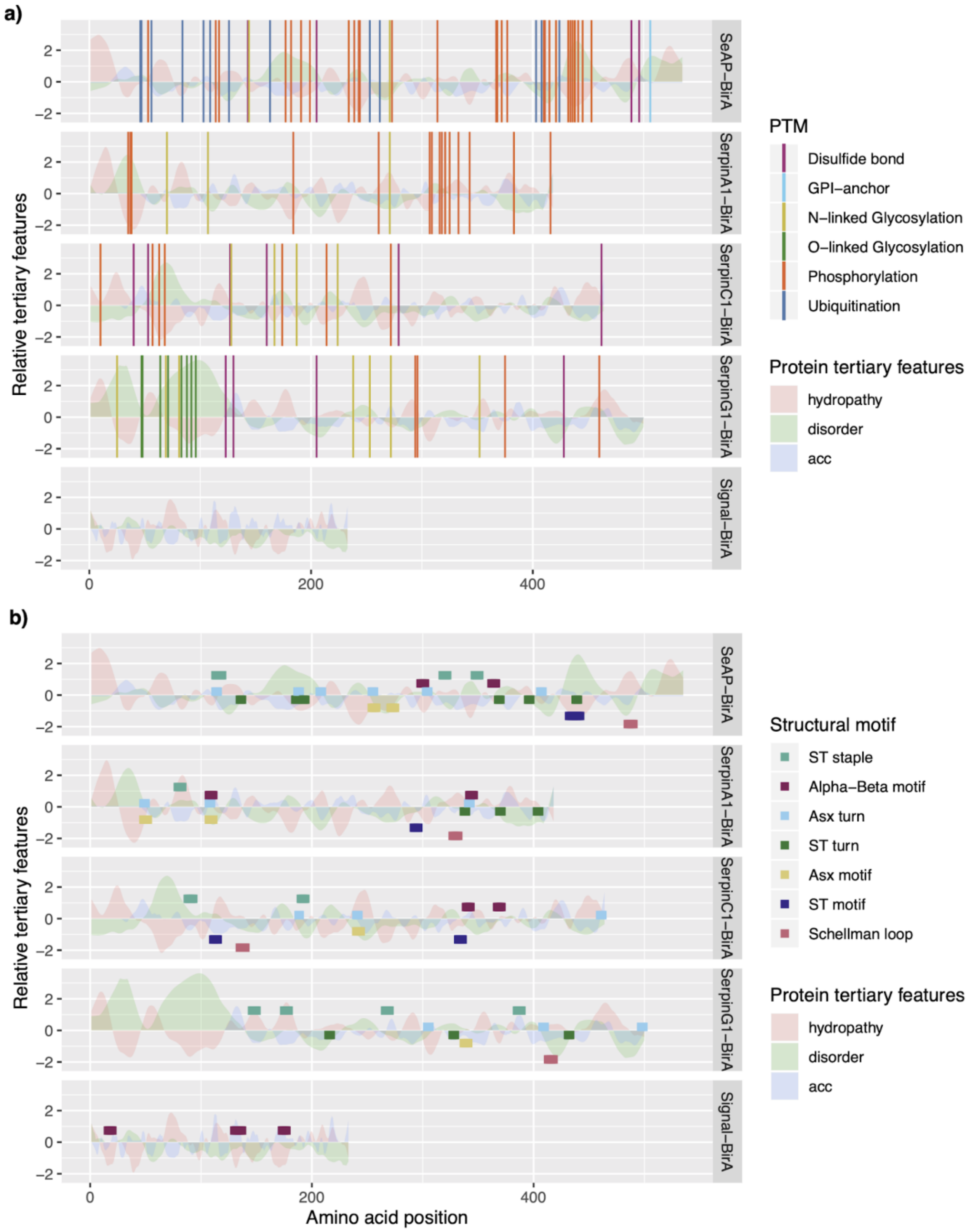


**Figure S4. The bait proteins show diversity in their PTM and structural content.** Dots and lines represent known PTM sites and structural motifs in panel (a) and (b) respectively. The hills and valleys indicate protein tertiary features. Note that solvent accessibility and structural motifs are only available for regions covered by the PDB structure, whereas predicted features such as protein hydrophobicity and disorder are available for the entire protein.

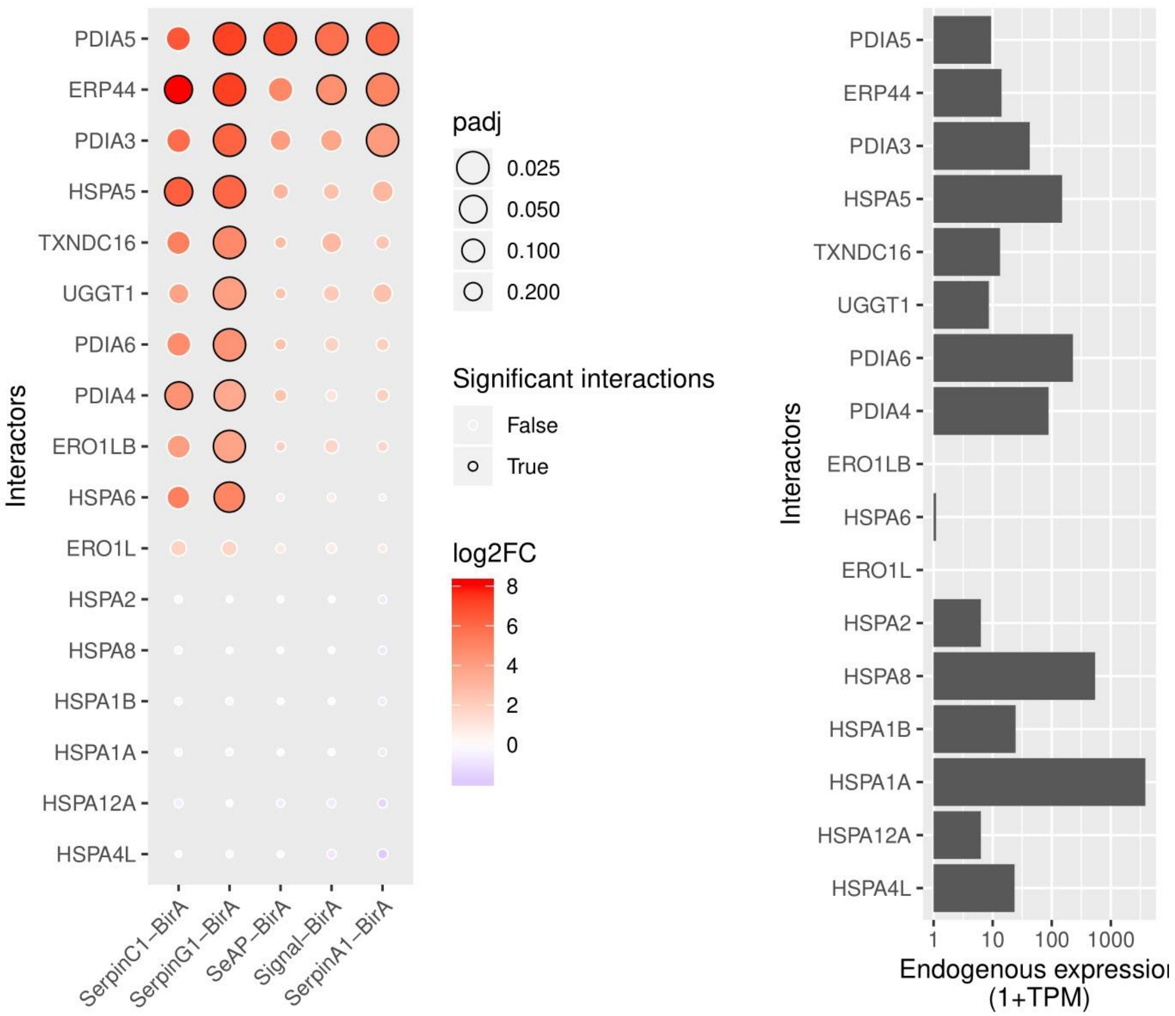


**Figure S5. Several proteins from the PDI family and molecular chaperones were selected from the significant interactors.** Panel (a) and (b) show their interaction intensity with the model proteins and their endogenous expression levels in HEK293 cells (Uhlén et al., 2015) respectively.
